# Tissue specificity of *in vitro* drug sensitivity

**DOI:** 10.1101/085357

**Authors:** Fupan Yao, Seyed Ali Madani Tonekaboni, Zhaleh Safikhani, Petr Smirnov, Nehme El-Hachem, Mark Freeman, Venkata Satya Kumar Manem, Benjamin Haibe-Kains

**Affiliations:** Princess Margaret Cancer Centre, Toronto, Ontario M5G 1L7, Canada; Department of Medical Biophysics, University of Toronto, Toronto, Ontario M5G 1L7, Canada; Integrative systems biology, Institut de Recherches Cliniques de Montréal, Montreal, Quebec, Canada; Department of Medicine, University of Montreal, Montréal, Quebec, Canada; Department of Computer Science, University of Toronto, Toronto, Ontario M5T 3A1, Canada; Ontario Institute of Cancer Research, Toronto, Ontario M5G 1L7, Canada

## Abstract

Research in oncology traditionally focuses on specific tissue type from which the cancer develops. However, advances in high-throughput molecular profiling technologies have enabled the comprehensive characterization of molecular aberrations in multiple cancer types. It was hoped that these large-scale datasets would provide the foundation for a paradigm shift in oncology which would see tumors being classified by their molecular profiles rather than tissue types, but tumors with similar genomic aberrations may respond differently to targeted therapies depending on their tissue of origin. There is therefore a need to reassess the potential association between pharmacological response and tissue of origin for therapeutic drugs, and to test how these associations translate from preclinical to clinical settings.

In this paper, we investigate the tissue specificity of drug sensitivities in large-scale pharmacological studies and compare these associations to those found in clinical trial descriptions. Our meta-analysis of the four largest *in vitro* drug screening datasets indicates that tissue of origin is strongly associated with drug response. We identify novel tissue-drug associations, which may present exciting new avenues for drug repurposing. One caveat is that the vast majority of the significant associations found in preclinical settings do not concur with clinical observations. Accordingly, our results call for more testing to find the root cause of the discrepancies between preclinical and clinical observations.

## INTRODUCTION

Large projects, such as The Cancer Genome Atlas [1] and the International Cancer Genome Consortium [2], have enabled the comprehensive characterization of molecular aberrations in multiple cancer types. The collection of mutations, copy number variations, gene expressions and other features enabled molecularly-based patient stratification across diverse tumor types, potentially creating a shift from the traditional classification based on tissue type [3–5]. However, tumors with similar genomic aberrations may respond differently to cytotoxic and targeted therapies, suggesting that tissue-of-origin is unlikely to be supplanted by molecular stratifications [6].

Testing drug potency in large populations of patients suffering different cancer types is an expensive and lengthy process [7]. Cancer cell lines provide a safe and cost-efficient methodology to measure drug response in multiple cancer types [8]. However, the translation of these preclinical findings in animal studies [9, 10] and in clinical settings [11] is complex, as cancer cell lines substantially differ from the patient tumors they originate from [12, 13]. This discrepancy has several causes. Repeatedly culturing cell lines allows for the potential acquisition of genomic aberrations, causing the cell lines to diverge from their initial samples [14]. In addition, mislabeling, simple clerical mistakes in cell line annotations, and cross-contamination can also cause additional skewing of drug screening results [15–17]. Despite these drawbacks, cell lines are the only model systems currently enabling high-throughput drug screening and will therefore remain models of choice for drug development and biomarker discovery [18–23].

In a recent paper investigating a pharmacogenomic dataset of 59 cell lines (NCI60), Jaeger et al. observed that drugs designed for specific tissue types, such as lapatinib for breast cancer, had similar activity across all tested tissue types, rather than a unique sensitivity pattern for the targeted tissue type [24]. Despite the small number of cell lines in NCI60, the authors concluded that cancer-specific drugs do not show higher efficacies in cell lines representing the tissue of interest, raising doubts about the relevance of *in vitro* screening for drug discovery and repurposing. If the results of this seminal study generalize to larger panel of cell lines, this would call for more curation of established cell lines to verify their tissues of origin, and for the generation of new cell lines or organoids freshly derived from patients as better models for high-throughput drug screening [10, 25, 26].

The recent release of multiple large-scale pharmacogenomic datasets enables analysis of sensitivities of thousands of cell lines to hundreds of drugs [18–21, 23]. Subsequent evaluation of these datasets, however, found only moderate inter-laboratory concordance in the drug response phenotypes [20, 27–30], highlighting the need for meta-analysis of these valuable, yet complex studies [31]. Such meta-analysis is hindered by the lack of standardization in cell line and drug identifiers. We addressed this issue by developing the *PharmacoGx* platform, which provides a computational system to allow for unified processing of pharmacogenomic datasets curated with standard cell line and drug identifiers [32].

In this work, we leveraged *PharmacoGx* to robustly assess the specificity of 727 experimental and approved drugs to 1527 unique cancer cell lines originating from 35 different tissue types. We then compared the significant drug-tissue associations identified *in vitro* to clinical observations. Our meta-analysis results indicate that tissue of origin is strongly predictive of drug response *in vitro.* However we found that, except for a few drugs, the vast majority of these preclinical associations did not concur with results from clinical trials, calling for further investigations of the relevance of cancer cell lines for drug sensitivities in specific tissue types.

## METHODS

The overall analysis design is represented in Figure 1.

**Figure 1:**
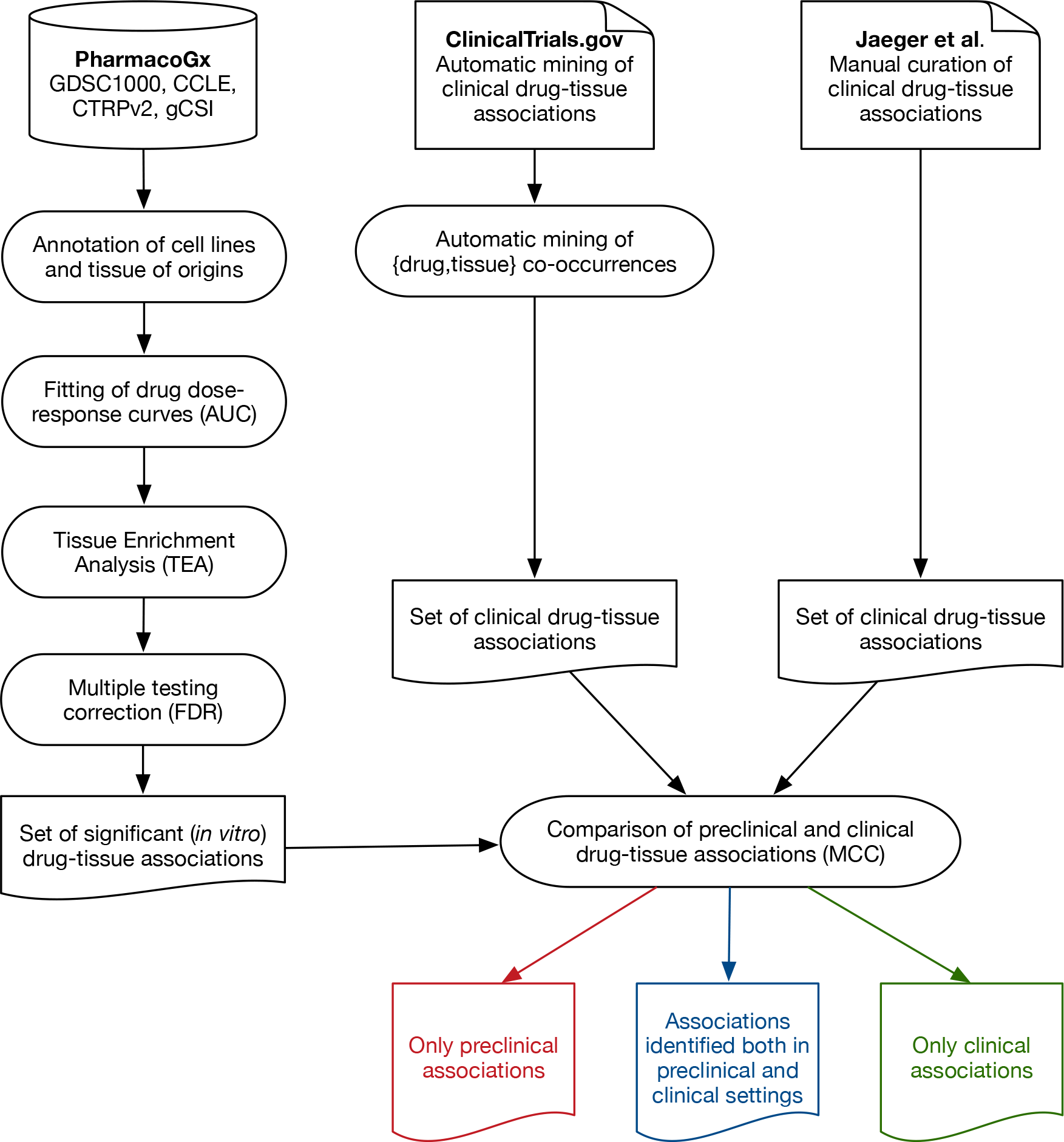
Schematic representation of data input and analysis pipeline.

### Pharmacogenomic datasets

We curated the four largest pharmacogenomic datasets within our *PharmacoGx* platform [32]: The Cancer Cell Line Encyclopedia (CCLE) [18], the Genomics of Drug Sensitivity in Cancer (GDSC1000) [19, 23, 33], the Cancer Therapy Response Portal (CTRPv2) [21, 34], and the Genentech Cell Line Screening Initiative (gCSl) [20] (Table 1). Cell lines were annotated using the Cellosaurus annotation database [35], while drugs were annotated using SMILES structures [36], PubChem ID [37], and InChiKeys [38]. All curated data were stored as *PharmacoSet* objects with our *PharmacoGx* platform (version 1.3.4) [32].

**Table 1:** **Characteristics of the pharmacogenomic datasets.**

|  | **CCLE** | **GDSC1000** | **CTRPv2** | **gCSI** |
| --- | --- | --- | --- | --- |
| # of cell lines | 1061 | 1124 | 887 | 410 |
| # of tissue types | 24 | 36 | 23 | 23 |
| # of drugs | 24 | 251 | 545 | 16 |
| # of drug dose-response curves | 11,670 | 225,480 | 395,263 | 6,455 |
| Pharmacological Assay | CellTiter Glo | Syto60 | CellTiter Glo | CellTiter Glo |
| Data source | broadinstitute.org/ccle/ | cancerrxgene.org/ | broadinstitute.org/ctrp/ | research-pub.gene.com/gCSI-cell-line-data |
| Reference | [18] | [23] | [21] | [20] |

### Tissue of origin of cancer cell lines

We used the Catalog of Somatic Mutations in Cancer (COSMIC) nomenclature to consistently annotate cancer cell lines with their tissue of origin [39]. Tissues with less than 5 cancer cell lines were removed in each dataset to ensure sufficient sample number for subsequent analyses.

### Drug sensitivity

To ensure consistent evaluation of drug sensitivity, we used our *PharmacoGx* platform to reprocess the drug dose-response curves in our compendium of pharmacogenomic datasets [32]. All dose-response curves were fitted to the equation 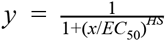, where *y* = 0 denotes death of all cancer cells within a sample, *y* = *y*(0) = 1 denotes no effect of the drug dose on the cancer cell sample, *EC*_50_ is the concentration at which viability is reduced to half of the viability observed in the presence of an arbitrarily large concentration of drug, and Hill Slope (*HS*) is a parameter describing the cooperativity of binding. *HS* < 1 denotes negative binding cooperativity, *HS* = 1 denotes noncooperative binding, and *HS* > 1 denotes positive binding cooperativity. The parameters of the curves were fitted using the least squares optimization framework. We used the area above the dose-response curves (AUC ∈ [0, 1]) to quantify drug sensitivity across cell lines, as AUC is always defined (as opposed to IC_50_) and combines potency and efficacy of a drug into a single parameter [40]. In this work, high AUC is indicative of sensitivity to a given drug.

### Tissue specificity of drug sensitivity

*Identification of drug-tissue associations using enrichment analysis.* For each drug, cell lines were first ranked based on their drug sensitivity (AUC) in each dataset separately. We then adapted the gene set enrichment analysis [41] implemented in the *piano* package [42] to test whether this ranked list is enriched in sensitive cell lines belonging to a specific tissue type (Supplementary Figure 1). Our tissue enrichment analysis (TEA) therefore allowed us to compute the significance of the association between each tissue and drug sensitivity using 10,000 cell line permutations in the tissue set. TEA was performed for each drug separately.

*Meta-analysis of drug-tissue associations*: Applying TEA to each dataset generates a set of p-values for each drug-tissue association. These p-values were combined using the weighted Z method [43] implemented in the *combine.test* function of our *survcomp* package [44]. Weights were defined as the number of cell lines in a given tissue type in each dataset from which the p-value has been computed. These combined p-values were subsequently adjusted using the false discovery rate (FDR) procedure [45] for all drugs.

### Clinical drug-tissue associations

To extract known drug-tissue association from the literature, we downloaded the list of clinical trials and associated metadata from ClinicalTrials.gov (last updated October 10th, 2016) and parsed these data using the XML R package (version 3.98-1.4). We then mined these clinical data to identify drug and tissue names. The co-occurrence of a drug with a tissue type was considered to constitute a drug-tissue association supported by clinical evidence. We also complemented our set of clinical drug-tissue associations by including the curation done by Jaeger et al. (Additional Files 1 and 2 in [24]).

### Comparison of drug-tissue associations between preclinical and clinical settings

To test whether drug-tissue associations observed in clinical trials were recapitulated *in vitro,* we compared the sets of preclinical and clinical associations by restricting our analysis to the associations tested in our meta-analysis of the pharmacogenomic data. We used Cytoscape version 3.3.0 [46] to visualize the associations observed in preclinical, clinical settings, or both as a network with colored edges in a Circos plot [47]. The Matthew correlation coefficient (MCC) [48] was used to quantify the strength of association between preclinical and clinical drug-tissue associations, and the significance was computed using a permutation test as implemented in the *PharmacoGx* R package (version 1.3.4) [32].

### Research replicability

This study complies with the standards of research reproducibility published by Sandve et al. [49]. The datasets are freely available through our *PharmacoGx* platform [32]. The code to replicate the analysis results, figures and tables is open-access and available on GitHub (github.com/bhklab/DrugTissue). In addition, we have set up a Docker virtual environment [50] is available online with all required R packages and tools pre-installed to facilitate replication of the study results. A detailed description of the software environment and the main steps to replicate the figures and tables is provided in Supplementary Information.

## RESULTS

Given the increasingly prominent use of high-throughput *in vitro* testing in biomedical research, we sought to test whether cancer cell lines originating from specific tissues responded differently to a large set of cytotoxic and targeted therapies. We therefore analyzed the four largest pharmacogenomic datasets published to date, namely CCLE, GDSC1000, CTRPv2 and gCSI (Table 1). These datasets contain 24 tissue types represented by at least 5 cell lines across all datasets (Figure 2A). Importantly, our curation [27, 30, 32] revealed that these studies investigated many identical cell lines and drugs, including 303 cell lines and 4 drugs—erlotinib, lapatinib, paclitaxel and crizotinib—screened in all four datasets (Figure 2B; Supplementary File 1).

**Figure 2:**
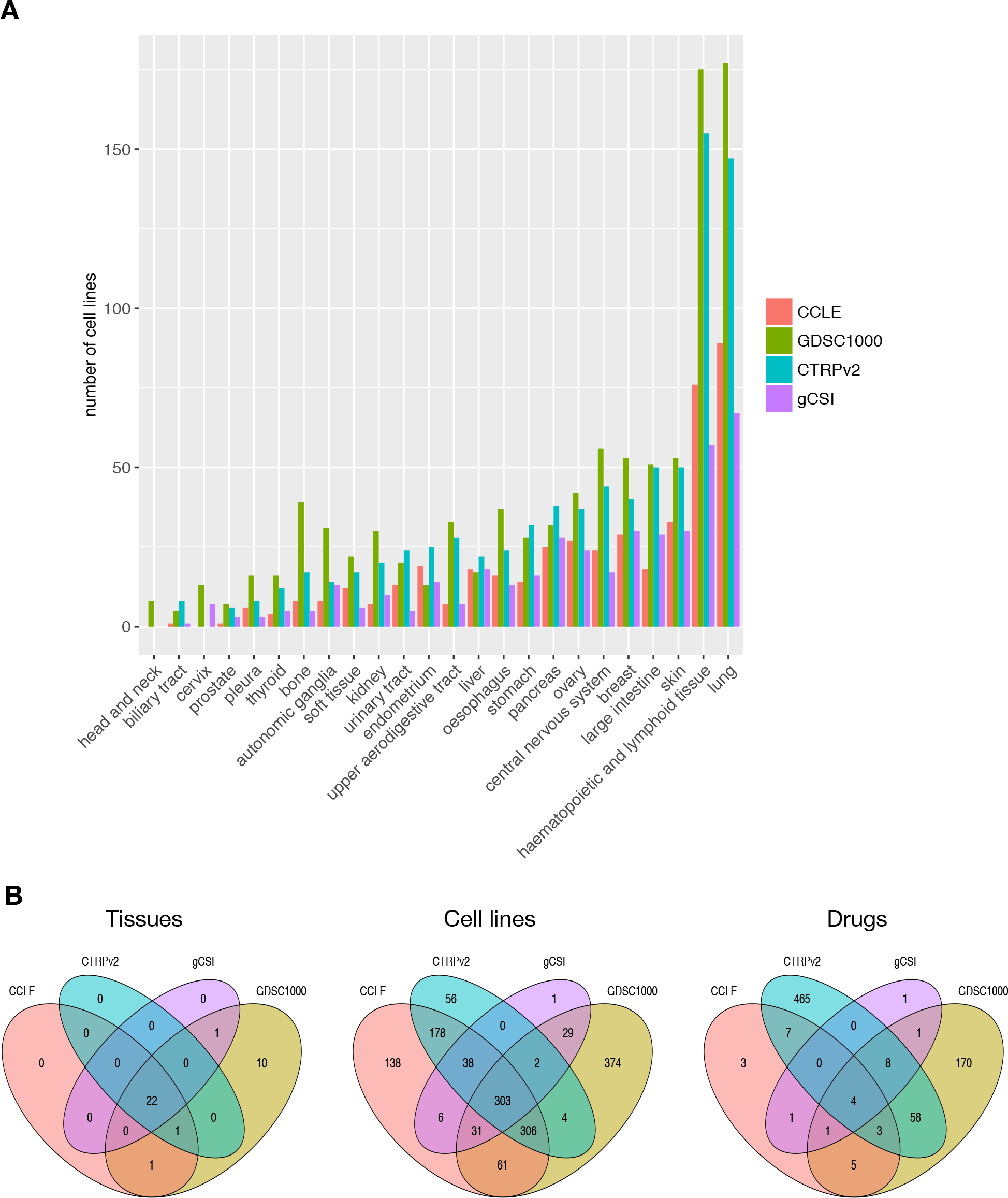
Composition and overlap of our compendium of pharmacogenomic datasets. (**A**) Number of cell lines representing each tissue type with respect to their source dataset. Tissue types represented by less than 5 cell lines in a given dataset were removed for the dataset. (**B**) Overlap for drugs, cell lines and tissue types across datasets.

We leveraged our compendium of pharmacogenomic datasets to identify statistically significant drug-tissue associations *in vitro* using our Tissue Enrichment Analysis (TEA; Supplementary Figure 1). We reported the enrichment scores and associated p-values for the 16 drugs screened in at least 3 datasets in Supplementary File 2. Out of 722 drugs in our data compendium, we found that 631 (87%) of drugs yielded significantly higher sensitivities in at least one tissue type (FDR<5%; Supplementary File 3). Interestingly, 17-AAG, trametinib, afatinib, docetaxel yielded significantly enriched sensitivities in more than 5 tissue types (Figure 3A-D), while less than 10% of the drugs, such as betulinic acid and MS275, showed no dependency of sensitivity to tissues (Figure 3E,F).

**Figure 3:**
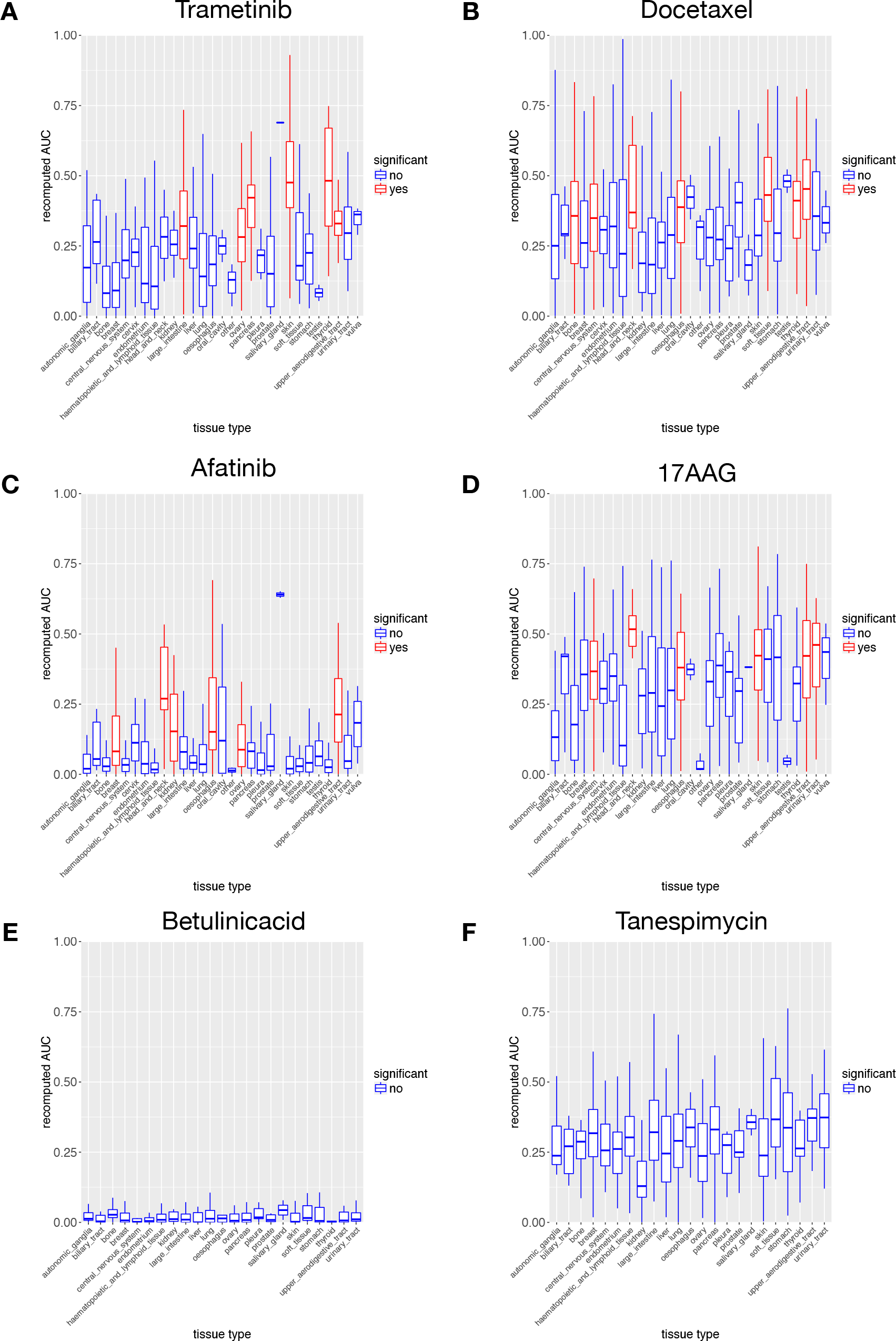
Distribution of drug sensitivities with respect to tissue types for drugs yielding the strongest(A-D) and weakest (E-F) tissue associations. Tissue enriched in drug sensitivities are highlighted in red.

Despite the large number of drugs with enriched sensitivity in at least one tissue type, only 6% of all the drug-tissue associations assessed in our study (1,025/17,371) were significant (FDR<5%). We investigated whether these significant associations were uniformly distributed across tissues types. There was no significant correlation between the number of drug-tissue associations and the number of cell lines in each tissue type (Spearman correlation coefficient ρ=0.05, p-value=0.80, which was expected because TEA controls for the size of tissue sets during the permutation testing procedure. We found that the pleura tissue and breast luminal B subtype of breast cancer have no drug-tissue associations (Figure 4A). Concurring with previous reports [51, 52], the “hematopoietic and lymphoid” tissue was significantly enriched in sensitive cell lines for more than 500 drugs (65%; Figure 4A), suggesting that these cell lines are highly sensitive to chemical perturbations. To examine the impact of this tissue on the overall analysis, TEA was rerun removing all the corresponding cell lines (217 unique cell lines). The number of significant drug-tissue associations proportionally decreased (6% v.s. 3% for the full analysis and without hematopoietic and lymphoid tissue, respectively), suggesting that the associations with other tissues are not drastically influenced by the presence of the hematopoietic and lymphoid tissue.

**Figure 4:**
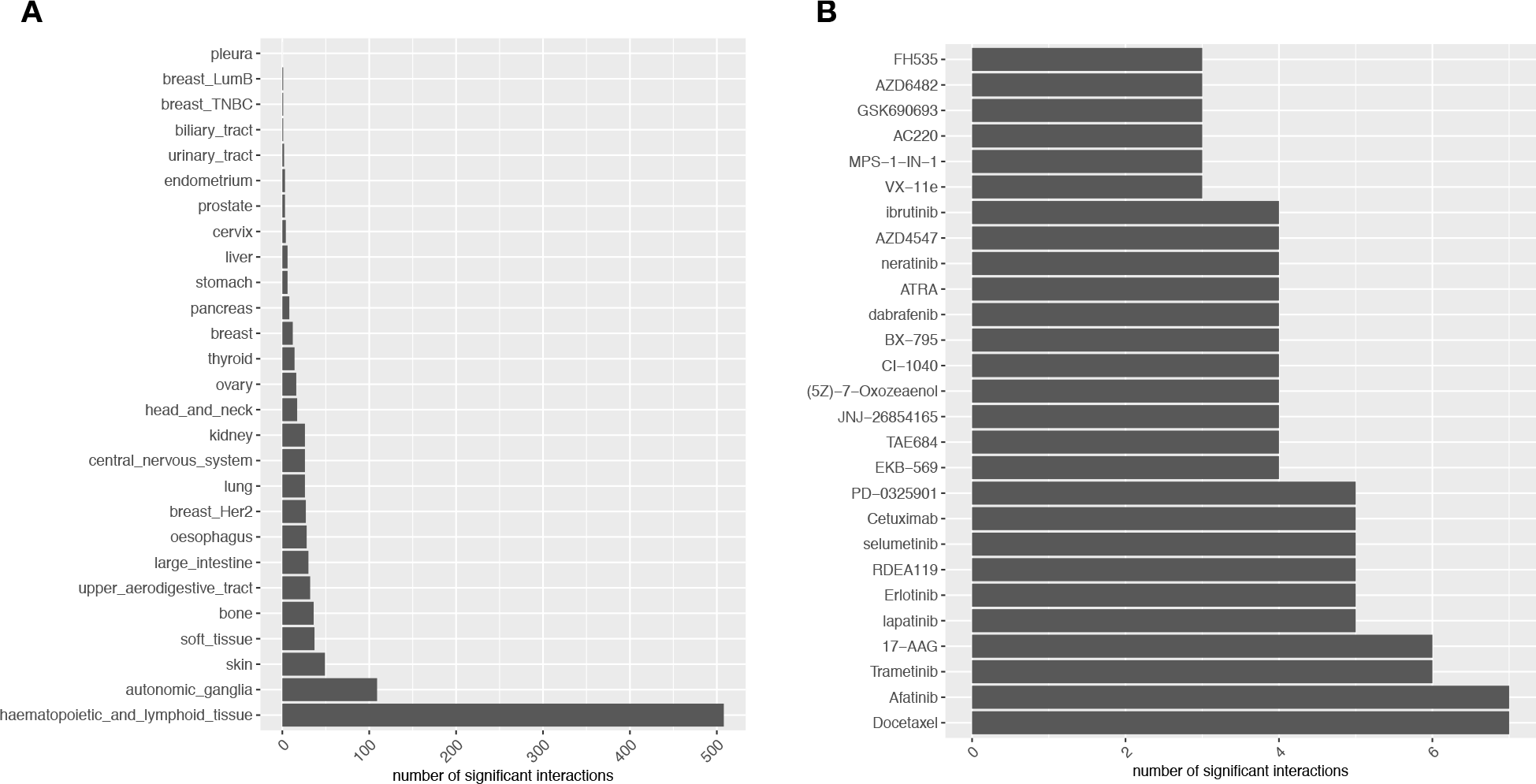
Distribution of *in vitro* drug-tissue associations. (**A**) Number of significantly associated drugs for each tissue type in our compendium; (**B**) Number of significantly associated tissues for drugs with 3 or more associations.

Although our meta-analysis leverages the four largest pharmacogenomic studies published to date, these datasets vary in terms of the number of drug dose-response curves actually measured (Table 1). We therefore assessed which dataset contributed the most to the discovery of statistically significant *in vitro* drug-tissue associations. As expected, the two largest datasets, namely GDSC1000 and CTRPv2, contributed several times more associations than gCSI and CCLE (Figure 5). Interestingly, more than half of significant drug-tissue associations are “private”, i.e., they were found in one dataset but were not consistent across datasets. On the contrary, a substantial proportion of associations were not significant in each individual datasets but were selected during the meta-analysis phase based on their consistent trend to significance (Figure 5). These results support the benefit of combining multiple pharmacogenomic datasets in a meta-analysis framework.

**Figure 5:**
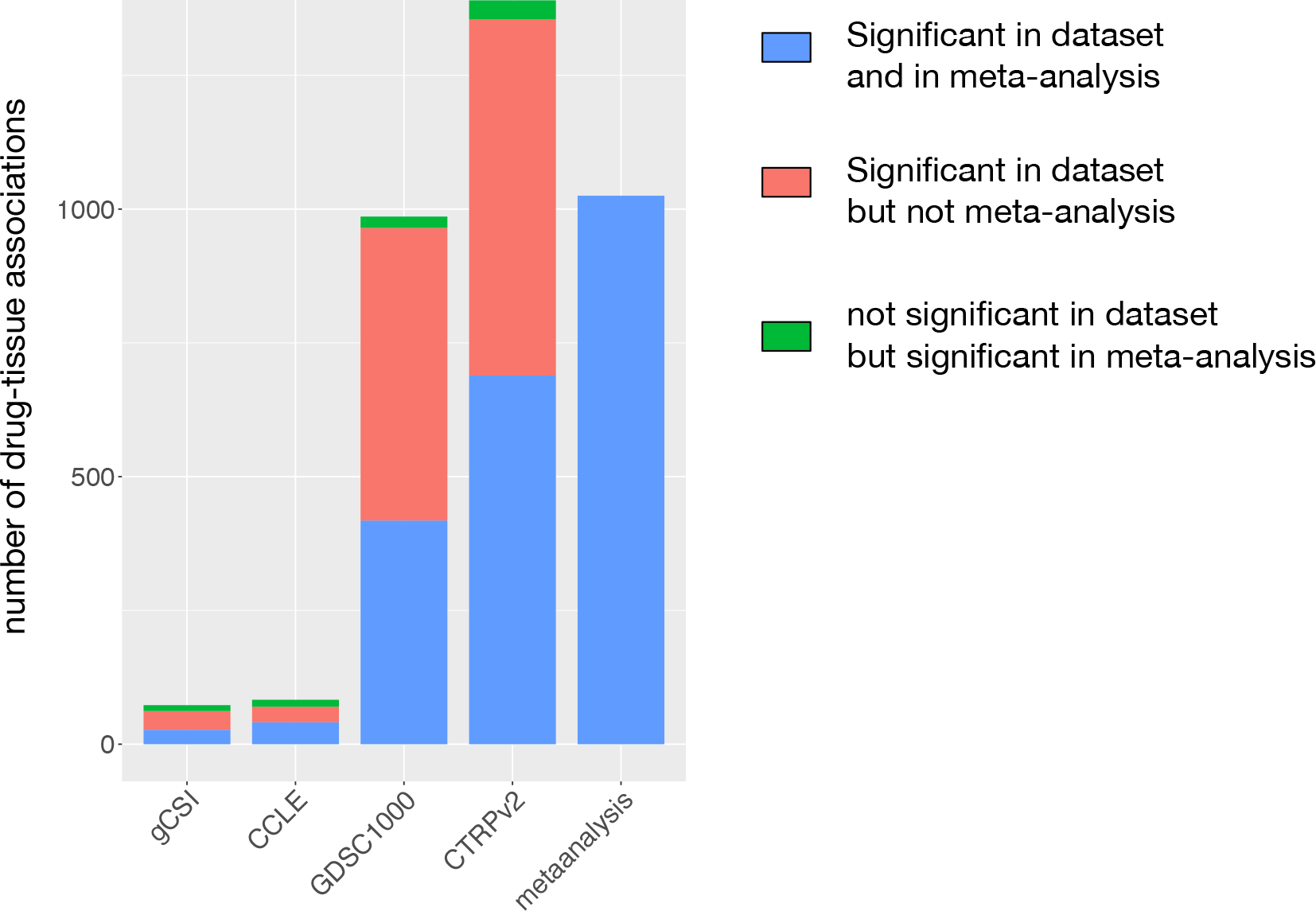
Number of drug-tissue associations in each pharmacogenomic dataset and meta-analysis. The associations that are significant in a dataset and also in the meta-analysis are colored in blue. The associations found significant in a dataset but not selected after meta-analysis are colored in red. The associations found non-significant in a dataset but ended up selected after meta-analysis are colored in green.

We further investigated the impact of sample size, that is the number of cell lines screened for drug sensitivity, on the TEA. When restricting our analysis to the NCI60 cell line panel (59 cell lines from 9 tissue types) and the 86 drugs in common with our compendium, we found less significant drug-tissue associations (16 v.s. 80 for NCI60 and our compendium, respectively; Supplementary Figure 2), indicating that the number cell lines in NCI60 is not sufficient to identify most of the drug-tissue associations. Analyzing the full set of 50,978 drugs in NCI60, we also found a substantially lower proportion of drug-tissue associations to be significant for the drugs and 9 tissue types present in our NCI60 PSet (6% v.s. 0.22% with FDR<5% for our compendium and NCI60, respectively; Supplementary Figure 3; Supplementary Files 3 and 4).

Given the significant tissue specificity of most drugs *in vitro*, we sought to assess whether these associations were consistent with clinical observations regarding the efficacy of drugs in specific tissue types. We therefore mined a large set of clinical trials to extract co-occurrences of tissues and drugs in the trial descriptions published in ClinicalTrials.gov. Distribution of co-occurrences for each drug with tissue types is shown in Figure 6A. We found 535 potential drug-tissue associations tested in clinical settings. Despite the large number of *in vitro* and clinical associations, the overlap was small (MCC=-0.004, p=0.99; Figure 6B). We confirmed the lack of global concordance between preclinical and clinical drug-tissue associations using the manual curation of Jaeger et al. (Figure 6C). Interestingly, we observed that afatinib and lapatinib, two of the drugs with the most number of tissue associations *in vitro*, are part of the small set of conserved drug-tissue associations (Table 2; Figure 7), suggesting that the strongest *in vitro* associations can be translated into clinical setting. However, we consider such a finding to be weak at best as this represents a very small set of associations.

**Figure 6:**
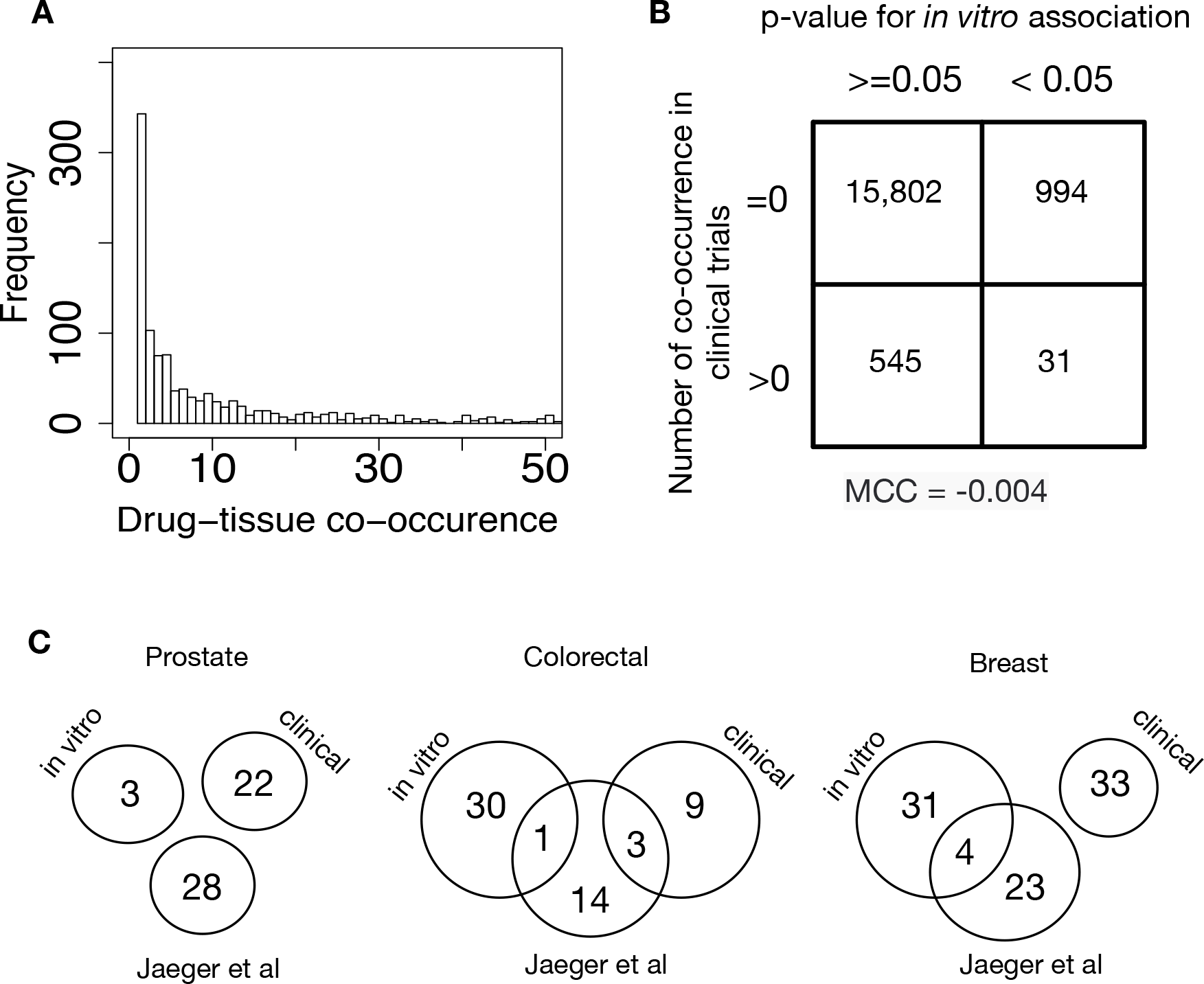
Comparison of preclinical and clinical drug-tissue associations. (**A**) Distribution of (drug, tissue) co-occurrences from records in ClinicalTrials.gov. **(B**) Contingency table for the drug-tissue association *in vitro* and drug-tissue co-occurrence in clinical trials. (**C**) overlap of interactions found *in vitro* and in clinical data from Jaeger et al and clinical trials. (**D**) interaction graph showing clinical trial data against preclinical data: Red lines represent drug-tissue associations observed only *in vitro* (referred to as experimental). Pink lines indicate experimental relationships with no clinical application. Green lines indicate a clinical application not recognized in preclinical analysis. Blue lines indicate *in vitro* drug-tissue associations supported by clinical applications.

**Table 2:** **List of drug-tissue associations conserved across *in vitro* and clinical settings.**

| **Drug** | **Tissue** | **ClinicalTrials.gov**<br>**(# co-occurrences)** | **Jaeger et al** | ***in vitro* FDR** |
| --- | --- | --- | --- | --- |
| Erlotinib | Kidney | 19 | yes | 1.11e-05 |
| Cisplatin | Cervix | 12 | no | 2.95e-19 |
| Temsirolimus | Thyroid | 27 | yes | 0.02 |
| Afatinib | Breast | 23 | yes as Afatinib | 0.001 |
| Lapatinib | Kidney | 2 | no | 1.3e-5 |
| Lapatinib | Breast | 525 | yes | 0.0006 |
| Pazopanib | Liver | 5 | no | 0.007 |
| XL765 | Lung | 1 | no | 2.95e-19 |
| Dabrafenib | Skin | 1 | no | 4.24e-35 |

**Figure 7:**
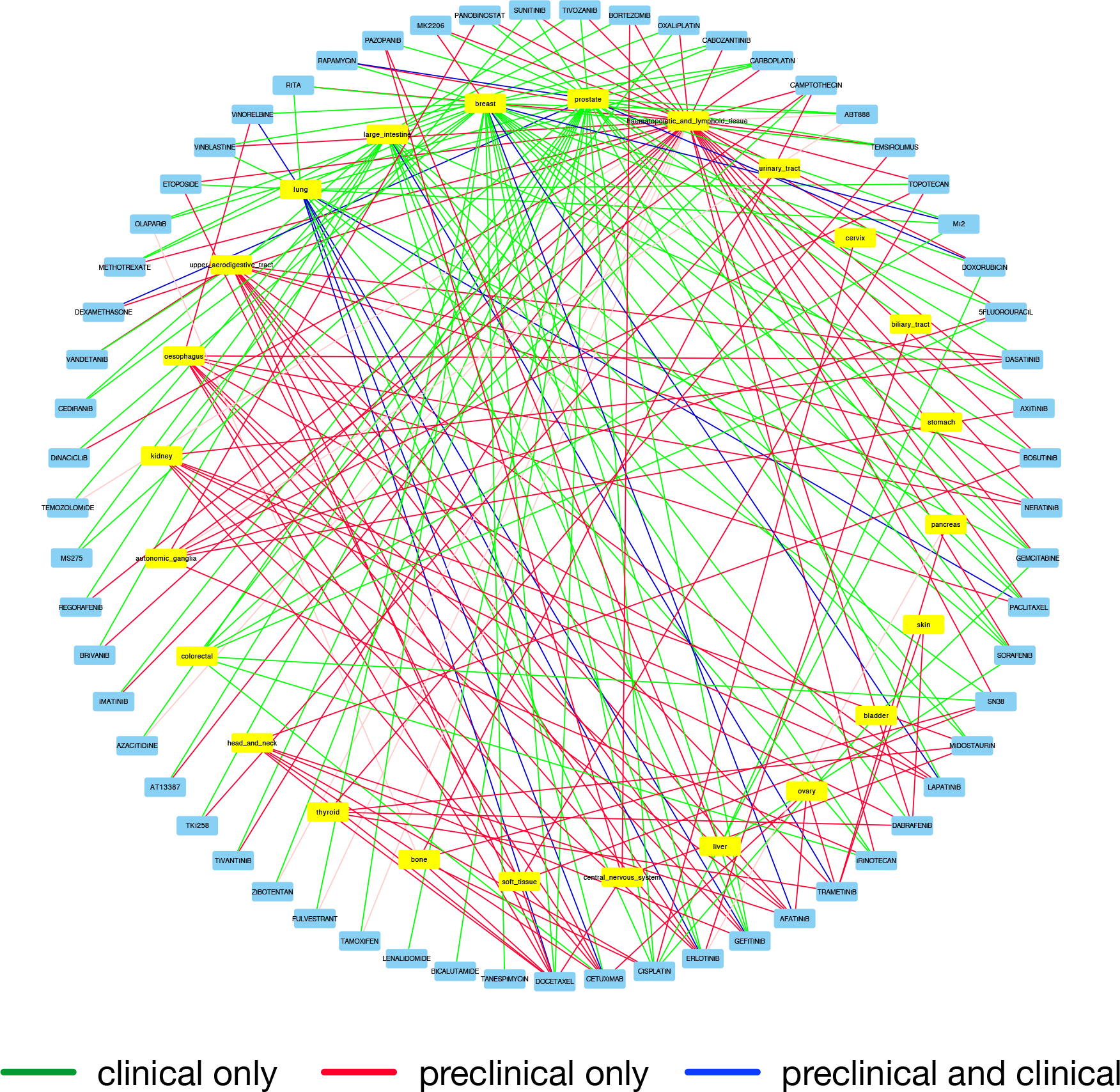
Circos plot representing the significant associations for drugs with clinical trial evidence. Blue and yellow boxes represent the drugs and tissues, respectively. Red lines represent drug-tissue associations observed only *in vitro* (referred to as experimental). Pink lines indicate experimental relationships with no clinical application. Green lines indicate a clinical application not recognized in preclinical analysis. Blue lines indicate *in vitro* drug-tissue associations supported by clinical applications.

## DISCUSSION

One of the main challenges in precision cancer medicine is to select drugs likely to yield response for each patient. Most of the current treatment regimens for cancer are based on the tissue of origin, as therapies are being designed for specific tissues [6, 53]. Recent high-throughput *in vitro* drug screening studies investigating large panels of cancer cell lines from multiple tissues [18–21, 23] provide us with a unique opportunity to assess the association between drug sensitivity and tissue types [53]. However, it remains unclear to which extent cancer cell lines originating from different tissue types respond differently to a variety of cytotoxic and targeted drugs [24, 28, 51, 52, 54, 55]. If these drug-tissue associations recapitulated the differential drug response across tissues observed in clinic, this would open a new avenue of research for tissue-based drug repurposing. In this study, we addressed these two issues in the largest meta-analysis of pan-cancer *in vitro* drug screening data published to date.

Our large compendium of drugs and cancer cell lines representing 25 tissue types, combined with our tissue enrichment analysis, allowed us to identify a large number of *in vitro* drug-tissue associations. Our results indicate that the vast majority of the drugs (87%) yield higher sensitivity in at least one tissue type. Our meta-analysis shed new light on the recent controversy regarding the tissue specificity of drug sensitivity screens, where studies reported substantial tissue-specific drug response [51, 52], and the contrary [24]. This controversy is partly due to the lack of consensus definition of tissue specificity. In our study and the previous work from Klijn et al. [51] and Gupta et al [52], tissue specificity is defined as an association between drug sensitivity and *any* tissue type, while Jaeger et al. [24] only considered associations with the tissues the drugs have been developed for. The latter set of associations is therefore a subset of all drug-tissue associations that can be identified *in vitro*. However, using the broader definition of tissue specificity, our results indicate that the small panel of 59 cell lines screened in the NCI60 dataset and used in Jaeger et al. [24] is not sufficient to identify most of the *in vitro* drug-tissue associations. Moreover, we also demonstrated that the meta-analysis framework used in this study, allowed us not only to identify more drug-tissue associations, but also to discard the drug-tissue associations that were not consistent across multiple datasets, increasing the robustness of our results.

While tissue-specificity of drug sensitivity *in vitro* is relevant for drug development in the preclinical setting, its translational potential in clinical setting remains unclear. In this regard, the study from Jaeger et al. was seminal as the authors compared *in vitro* drug response patterns to clinical observations in breast, colorectal and prostate cancers and found no concordance [24]. Given that our results indicate strong tissue specificity of *in vitro* drug sensitivity, we tested the concordance of preclinical and clinical observations in more than 23 tissue types. Although we found that the drugs showing the strongest tissue-specificity (afatinib and lapatinib) were more likely to be tested in clinical trials with their associated tissue types than the rest of the drugs, we observed no global overlap in drug-tissue associations between the preclinical and clinical settings. Concurring with Jaeger et al. [24], our results call into question the translational potential of the *in vitro* results.

We have come to recognize that cancer cell lines do not fully recapitulate the molecular features of patient tumors they originate from [11, 56], which may hinder the translation of ***in vitro*** drug development to clinical settings [55, 57–60]. It is hoped that large panels of cancer cell lines will enable faithful representation of the molecular diversity observed in patient tumours [18, 19, 23]. However, recent studies identified cell lines exhibiting molecular phenotypes that are not observed in patients [12, 13], casting doubts on the relevance of these model systems for biological investigation and drug screening. Another fundamental problem in cancer cell line studies is the lack of a standard nomenclature to uniquely annotate cell lines to their tissue of origin [16, 61, 62], even though ontologies are under active development [35, 63]. Lastly, cancer cell lines lack the tumor microenvironment, which has recently been shown to have a substantial effect on drug response and resistance [64, 65]. Patient-derived organoids and xenografts are now new models of choice for drug screening and their usage might alleviate the current limitations of cancer cell lines [9, 10, 25]. These are key factors that are likely to contribute to the discrepancy between preclinical and clinical observations highlighted in this study. Although our meta-analysis provides the largest repository of *in vitro* drug-tissue associations to date, our results call for further investigations with the aim to improve the translational potential of cancer cell lines.

## CONCLUSION

Our meta-analysis of pan-cancer *in vitro* drug screening datasets indicates that most approved and experimental drugs exhibit tissue-specific sensitivities in large panel of cancer cell lines. However, it is equally clear that the preclinical results do not translate to clinical setting, as the vast majority of *in vitro* drug-tissue associations are not recapitulated in clinical trials. Our results suggest that additional research is required to improve the translational potential of cancer cell lines for drug screening.

## ACKNOWLEDGEMENTS

The authors would like to thank Drs. Samira Jaeger and Patrick Aloy for their constructive and insightful feedback on our study.

## AUTHOR CONTRIBUTIONS

F.Y, Z.S and S.A.A.T contributed equally to this work. F.Y, Z.S and S.A.A.T wrote the code, performed the analysis and interpreted the results. Z.S, P.S and M.F collected and fitted the drug dose-response curves. N.E-H curated the cell lines and drug annotations. F.Y and V.S.K.M performed the tissue enrichment analysis. B.H.K supervised the study. F.Y, S.A.A.T, V.S.K.M wrote the first version of the manuscript.

## FUNDING

This study was conducted with the support of the Canadian Cancer Research Society and the Ontario Institute for Cancer Research through funding provided by the Government of Ontario. S.A.A.T was supported by a Connaught International Scholarship. Z.S was supported by the Cancer Research Society (Canada). N.E-H was supported by the Ministry of Economic Development, Employment and Infrastructure and the Ministry of Innovation of the Government of Ontario. B.H.K was supported by the Gattuso-Slaight Personalized Cancer Medicine Fund at Princess Margaret Cancer Centre and the Canadian Institutes of Health Research.

## COMPETING FINANCIAL INTERESTS

The authors declare no competing financial interests.

## SUPPLEMENTARY FIGURES

**Supplementary Figure 1:**
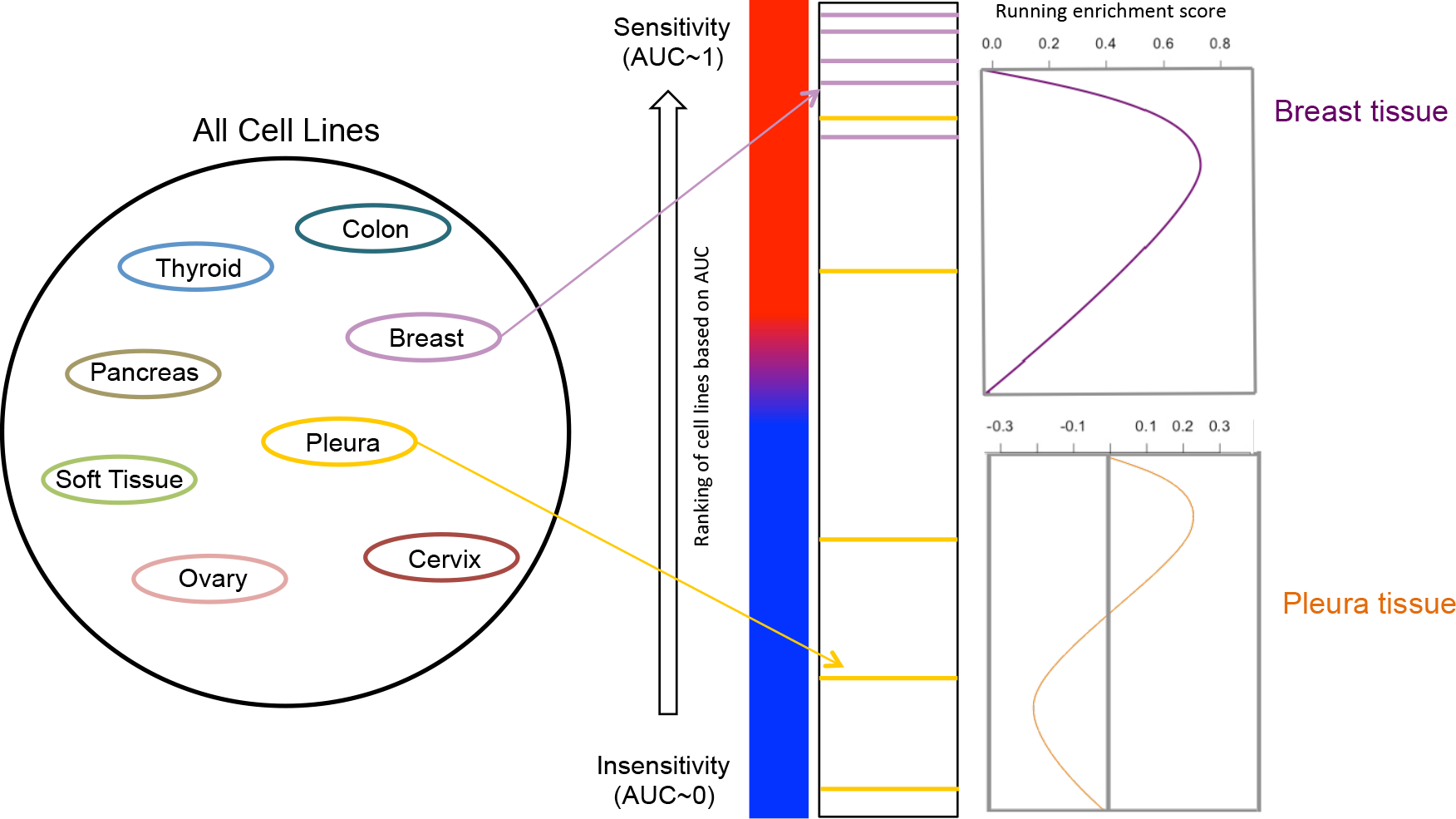
Schematic describing our tissue enrichment analysis (TEA). Our method is adapted from the gene set enrichment analysis (GSEA) where gene sets are replaced by sets of cell lines representing each tissue type. Breast is an example of strong drug-tissue association, as the breast cancer cell lines are all highly sensitivity (high AUC). On the contrary, the pleura tissue is not enriched in sensitive cell lines.

**Supplementary Figure 2:**
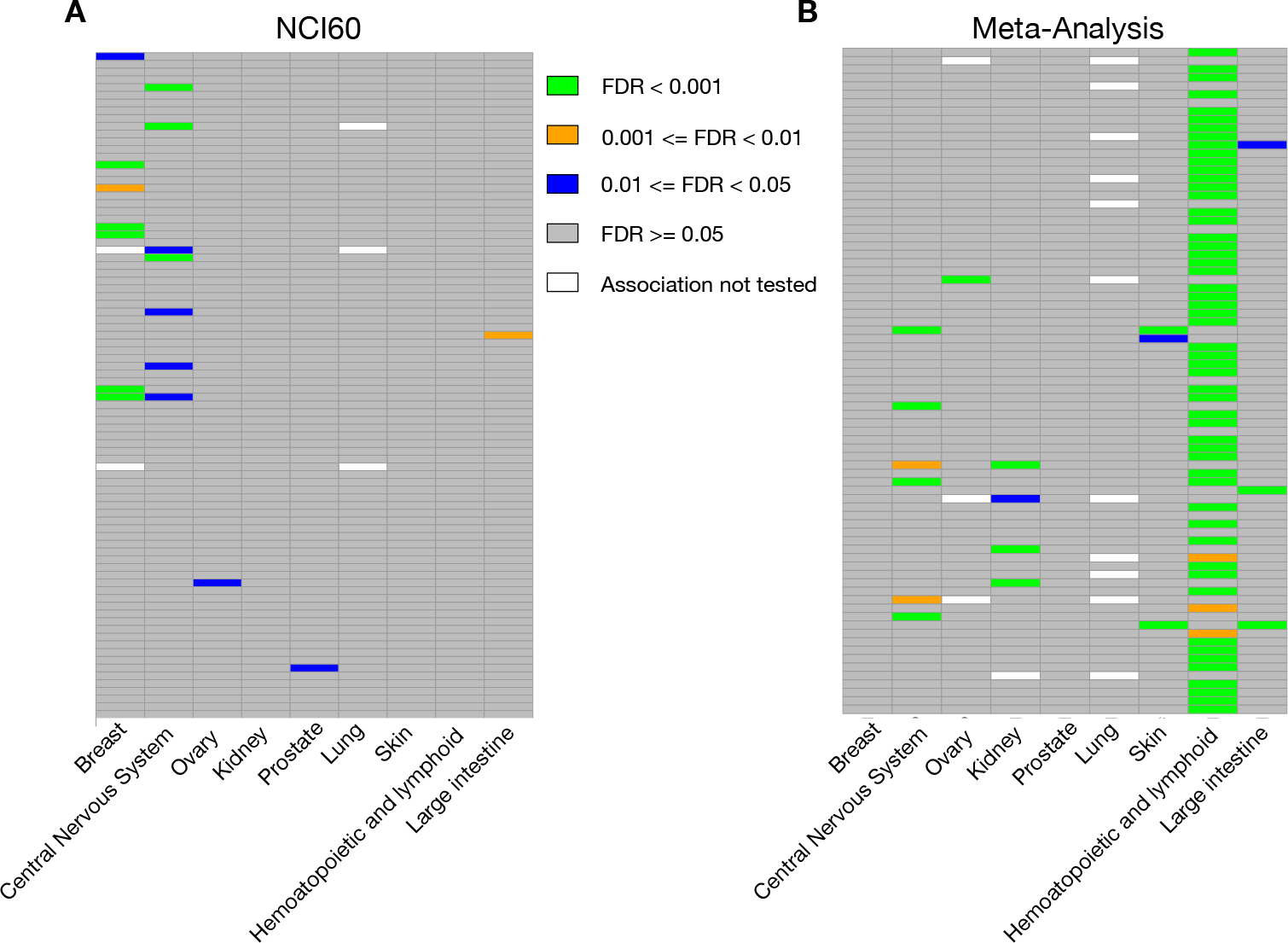
Heatmap showing the significance of the drug-tissue associations (FDR) for the drugs and tissue types in common between NCI60 and our compendium of datasets. (**A**) Tissue enrichment analysis (TEA) performed using the NCI60 data only. (**B**) TEA using our compendium of pharmacogenomic datasets. FDR < 0.001 in green; 0.001 ≤ FDR < 0.01 in orange; 0.01 ≤ FDR < 0.05 in blue; and FDR ≥ 0.05 in grey. Drug-tissue associations that were not tested due to lack of drug sensitivity data are represented white.

**Supplementary Figure 3: Heatmap showing the significance of the drug-tissue associations (FDR) for all the drugs and tissue types in the NCI60 dataset.** Only the drug with at least one association as determined by the tissue enrichment analysis (TEA) are represented. FDR < 0.001 in green; 0.001 ≤ FDR < 0.01 in orange; 0.01 ≤ FDR < 0.05 in blue; and FDR ≥ 0.05 in grey. Drug-tissue associations that were not tested due to lack of drug sensitivity data are represented white.

## SUPPLEMENTARY FILES

**Supplementary File 1: List of drugs in our compendium of pharmacogenomic datasets (CCLE, GDSC1000, CTRPv2, gCSI) and NCI60.**

**Supplementary File 2: Dataset-specific enrichment scores, associated p-values, and the false discovery rates for the drugs screened in at least 3 datasets.** Each sheet reports the results for a drug.

**Supplementary File 3: False discovery rates for all the the drug-tissue associations tested in our compendium of large pharmacogenomic datasets.** False discovery rates are computed from the combined p-values obtained from the tissue enrichment analysis performed in each dataset separately.

**Supplementary File 4: False discovery rates for all the the drug-tissue associations tested in the NCI60 dataset.** False discovery rates are computed from the p-values obtained from the tissue enrichment analysis.

## Supplementary Methods

### Full Reproducibility of the Analysis Results

We describe below how to fully reproduce the figures and tables reported in the paper

1. Set up the software environment
2. Run the R scripts
3. Generate figures

### Set up the software environment

We developed and tested our analysis pipeline using R running on linux and Mac OSX platforms. The code is freely available from GitHub https://github.com/bhklab/DrugTissue. The following is a copy of `sessionInfo()` from the development environment in R

~~~
R version 3.3.0 (2016-05-03)
Platform: x86_64-redhat-linux-gnu (64-bit)
Running under: CentOS Linux 7 (Core)
~~~

~~~
attached base packages:
[1] grid     stats    graphics    grDevices utils   datasets
[7] methods  base
~~~

~~~
other attached packages:
 [1] pvclust_2.0-0        vegan_2.4-0       lattice_0.20-33
 [4] permute_0.9-0        survcomp_1.22.0   prodlim_1.5.7
 [7] survival_2.39-4      piano_1.12.0      xlsx_0.5.7
[10] xlsxjars_0.6.1       rJava_0.9-8       RColorBrewer_1.1-2
[13] gplots_3.0.1         mgcv_1.8-12       nlme_3.1-128
[16] ggplot2_2.1.0        reshape2_1.4.1    VennDiagram_1.6.17
[19] futile.logger_1.4.1  PharmacoGx_1.1.6
~~~

~~~
loaded via a namespace (and not attached):
 [1] gtools_3.5.0       lsa_0.73.1           slam_0.1-35
 [4] sets_1.0-16        splines_3.3.0        colorspace_1.2-6
 [7] SnowballC_0.5.1    marray_1.50.0        sm_2.2-5.4
[10] magicaxis_1.9.4    BiocGenerics_0.18.0  lambda.r_1.1.7
[13] plyr_1.8.4         lava_1.4.3           stringr_1.0.0
[16] munsell_0.4.3      survivalROC_1.0.3    gtable_0.2.0
[19] caTools_1.17.1     labeling_0.3         Biobase_2.32.0
[22] parallel_3.3.0     Rcpp_0.12.5          KernSmooth_2.23-15
[25] relations_0.6-6    scales_0.4.0         limma_3.28.11
[28] gdata_2.17.0       rmeta_2.16           plotrix_3.6-2
[31] bootstrap_2015.2   digest_0.6.9         stringi_1.1.1
[34] SuppDists_1.1-9.2  tools_3.3.0          bitops_1.0-6
[37] magrittr_1.5       cluster_2.0.4        futile.options_1.0.0
[40] MASS_7.3-45        Matrix_1.2-6         downloader_0.4
[43] igraph_1.0.1
~~~

All these packages are available on CRANor Bioconductor All necessary packages have `library(<package>)` calls within the R scripts themselves, or the script assumes a previous script has been run and thus should have loaded necessary packages.

### Running R Scripts in Repository and Figure Generation

It is mandatory for all scripts to be run within one RStudio instance, as scripts will reference generated output variables from other script files

1. Load CTRPv2, CCLE, GDSC1000, and gCSI using the `downloadPSet()` functions in PharmacoGx
2. Run `GSEA_with_AUC.R` to get enrichment scores in one matrix variable called `combined1`
3. Load the XML clinicaltrials.gov reader from `drugResultGetter.R`, and run `wordmine.R` to generate the `wordmine` variable referenced in diagram generation
4. All figures should run independently with varying pdf generation in working directory set using `setWD()`
5. All figures can be generated with their respectively R file
6. The circos plot has been generated using Cytoscape for network visualization
7. Supplementary File 3 is generated at the end of `GSEA_with_AUC.R`
8. In `GSEA_with_AUC.R`, a boolean variable paper is present to recreate the TEA analysis without hematopoietic and lymphoid tissue mentioned in the paper

## 1 Acronyms

AUC: Area under the drug dose-response curve
CCLE: The Cancer Cell Line Encyclopedia initiated by the Broad Institute of MIT and Harvard
CGHub: The Cancer Genomics Hub from the University of California Santa Cruz and the US National Cancer Institute
COSMIC: Catalogue of Somatic Mutations in Cancer by the Wellcome Trust Sanger Institute
CTRPv2: The Cancer Therapeutics Portal version 2 initiated by the Broad Institute of MIT and Harvard
gCSI: The Genentech Cell Line Screening Initiative
GDSC1000: The Genomics of Drug Sensitivity in Cancer led by the Wellcome Trust Sanger Institute
MCC: Matthew Correlation Coefficient

